# Genomic analysis of *Mycobacterium tuberculosis* reveals complex etiology of tuberculosis in Vietnam including frequent introduction and transmission of Beijing lineage and positive selection for EsxW Beijing variant

**DOI:** 10.1101/092189

**Authors:** Kathryn E Holt, Paul McAdam, Phan Vuong Khac Thai, Dang Thi Minh Ha, Nguyen Ngoc Lan, Nguyen Huu Lan, Nguyen Thi Quynh Nhu, Nguyen Thuy Thuong Thuong, Guy Thwaites, David J Edwards, Kym Pham, Jeremy Farrar, Chiea Chuen Khor, Yik Ying Teo, Michael Inouye, Maxine Caws, Sarah J Dunstan

**Affiliations:** Centre for Systems Genomics, University of Melbourne, Parkville, Victoria 3010, Australia; Department of Biochemistry and Molecular Biology, Bio21 Molecular Science and Biotechnology Institute, University of Melbourne, Parkville, Victoria 3010, Australia; Pham Ngoc Thach Hospital for Tuberculosis and Lung Disease, Ho Chi Minh City, District 5, Viet Nam; Oxford University Clinical Research Unit, Ho Chi Minh City, District 5, Viet Nam; Centre for Tropical Medicine, Nuffield Department of Clinical Medicine, Oxford University, Oxford, UK.; Department of Pathology, University of Melbourne, Parkville, Victoria 3010, Australia; Genome Institute of Singapore, Singapore; Department of Statistics and Applied Probability, National University of Singapore, Singapore.; Saw Swee Hock School of Public Health, National University of Singapore, Singapore; School of BioSciences, University of Melbourne, Parkville, Victoria 3010, Australia; Liverpool School of Tropical Medicine, Liverpool, UK; Peter Doherty Institute for Infection and Immunity, University of Melbourne, Parkville, Victoria 3010, Australia

## Introduction

Tuberculosis (TB) is a leading cause of death from infectious disease and the global burden is now higher than at any point in history ^1,2^ Despite coordinated efforts to control TB transmission, the factors contributing to its successful spread remain poorly understood. Vietnam is identified as one of 30 high burden countries for TB and MDR-TB with an incidence of 137 TB cases per 100,000 individuals in 2015 ^2^ Recent phylogenomic analyses of the causative agent *Mycobacterium tuberculosis (Mtb*) in other high-prevalence regions have provided insights into the complex processes underlying TB transmission ^3–5^. Here we examine the transmission dynamics of *Mtb* isolated from TB patients in Ho Chi Minh City (HCMC), Vietnam via whole genome analysis of 1,635 isolates and comparison with 3,085 isolates from other locations. The genomic data reveal an underlying burden of disease caused by endemic *Mtb* Lineage 1 associated with activation of long-term latent infection, on top of which is overlaid a three-fold higher burden associated with introduction of exotic Lineage 2 and 4 *Mtb* strains. We identify frequent transfer of Beijing lineage *Mtb* into the country, which are associated with higher levels of transmission in this host population than endemic Lineage 1 *Mtb*. We identify a mutation in the secreted protein EsxW in Beijing strains that also appears to be under positive selection in other *Mtb* lineages, which could potentially contribute to the enhanced transmission of the Beijing lineage in Vietnamese and other host populations.

To characterize the diversity of *Mtb* circulating in HCMC, we sequenced the genomes of 1,635 isolates (**Supplementary Table 1**) obtained from 2,091 HIV uninfected, smear positive adults (≥18 years) commencing anti-TB therapy at district TB units (DTUs) in eight districts of HCMC between December 2008 and July 2011 (see **Methods**). A total of 73,718 high quality SNPs were identified and used to reconstruct a maximum likelihood (ML) phylogeny (**Fig. 1a**) and to assign lineages ^6^. Four major lineages (Lineages 1-4) were present within the study population. The majority of isolates (n=957, 59%) belonged to lineage 2.2.1, a subgroup of the Beijing lineage (2.2). Lineage 1 (Indo-Oceanic lineage; n=388, 23.7%) and Lineage 4 (EuroAmerican lineage; n=192, 11.7%) were also common. A single isolate belonged to Lineage 3 (East African-Indian lineage) and was excluded from further analysis. The distribution of lineages did not change during the 2.5 year period of the study (**Fig. 1b**), and was in agreement with previous genotyping (MIRU-VNTR, spoligotyping) studies in urban areas of Vietnam (≥50% Beijing lineage (2.2) and ~20% Lineage 1.1/EIA in Hanoi and HCMC, 1998-2009) ^7–11^. Known antimicrobial resistance mutations were detected in all lineages but were most frequent in Beijing sublineage 2.2.1 (**Table 1**), consistent with earlier reports from Vietnam ^7–9,11^.

**Figure 1.**
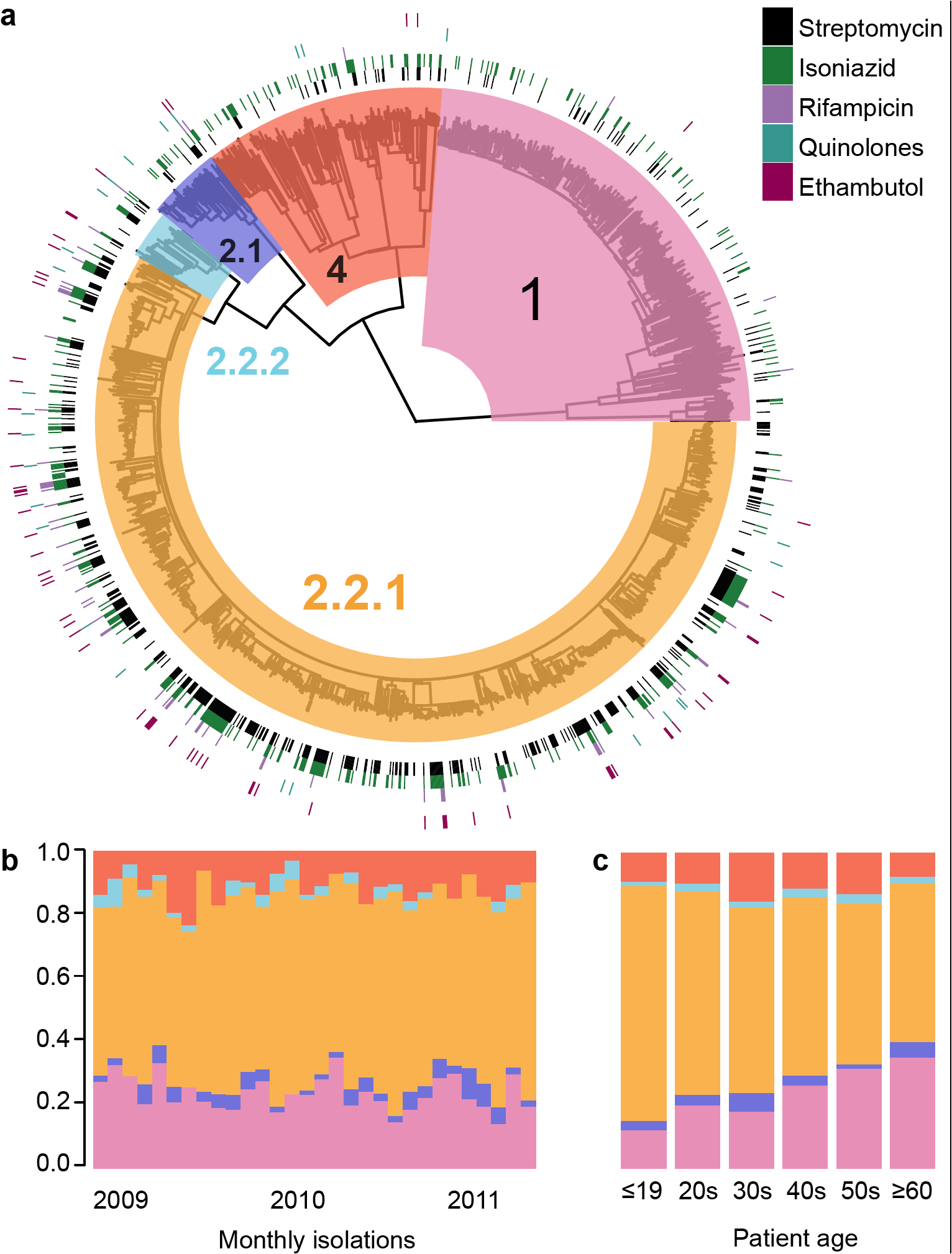
Circulating *M. tuberculosis* strains in HCMC are divided into multiple distinct lineages. **(a)** Maximum-likelihood phylogeny of HCMC isolates with backgrounds shaded by lineage. Exterior rings indicate presence of known antimicrobial resistance-associated mutations (coloured by drug, according to legend in top right). **(b)** Frequency distribution of lineages by month. **(c)** Frequency distribution of lineages by patient age group.

**Table 1.** Lineage characteristics for HCMC *M. tuberculosis* isolates, including known antimicrobial resistance mutations identified using Mykrobe Predictor.

|  | Lineage |  |  |  |  |
| --- | --- | --- | --- | --- | --- |
|  | 1 | 2.1 | 2.2.1 | 2.2.2 | 4 |
| **Gender** |  |  |  |  |  |
| Female | 82 (21.1%) | 9 (15.3%) | 265 (27.8%) | 10 (25.6%) | 56 (29.2%) |
| Male | 306 (78.9%) | 50 (84.7%) | 692 (72.3%) | 29 (74.4%) | 136 (70.8%) |
| **Antimicrobial** |  |  |  |  |  |
| Streptomycin | 48 (12.4%) | 10 (17.0%) | 426 (44.5%) | 12 (30.8%) | 30 (15.6%) |
| Isoniazid | 57 (14.7%) | 12 (20.3%) | 269 (28.1%) | 9 (23.1%) | 52 (27.1%) |
| Rifampicin | 3 (0.8%) | 2 (3.4%) | 58 (6.1%) | 2 (5.1%) | 1 (0.5%) |
| Quinolones | 1 (0.3%) | 3 (5.1%) | 18 (1.9%) | 2 (5.1%) | 2 (1.0%) |
| Ethambutol | 1 (0.3%) | 2 (3.4%) | 60 (6.3%) | 3 (7.7%) | 2 (1.0%) |

Whilst the majority of TB patients were male (74%, typical for TB studies in Vietnam and elsewhere ^8–10^), the Beijing lineage was significantly associated with TB in females (OR 1.28 [95% CI 1.01-1.62], p=0.043 using Fisher’s exact test; see **Table 1**). Beijing sublineage 2.2.1 was also significantly associated with younger people: its frequency declined with age, from 74% of cases in <20 year olds vs 50% in ≥60 year olds (p=0.0023 Fisher’s exact test, p=0.0024 linear trend test; **Fig. 1c**). In contrast, Lineage 1 was significantly associated with males (25% of male cases vs 19% of females, p=0.017) and increased with age regardless of gender, from 12% in <20 year olds vs 35% in ≥60 year olds (p=0.0007 Fisher’s exact test, p=0.0014 linear trend test; **Fig. 1c**). These data confirm that Beijing sublineage 2.2.1 is capable of infecting a wider range of hosts in the Vietnamese population, particularly among females and younger people, than is the endemic Lineage 1 ^7,10^.

Consequently, we hypothesised that Beijing lineage or sublineage 2.2.1 was also more transmissible than Lineage 1, and/or more capable of causing active disease in infected hosts, in the Vietnamese host population. We used the whole genome phylogeny to investigate this possibility in more detail, comparing several diversity metrics for each lineage (**Fig. 2**, **Supplementary Fig. 1**). Terminal branch lengths, which represent the upper bound of time since transmission for each *Mtb* case, were significantly shorter for Beijing sublineage 2.2.1 *Mtb* isolates (median 8 SNPs) than for non-Beijing lineage isolates (Lineage 1: median 53 SNPs, p < 1 × 10^−15^ using Kolmogorov-Smirnov test; Lineage 2.1: 30 SNPs, p < 1 × 10^−6^; Lineage 4: 17 SNPs, p < 1 × 10^−9^), and slightly shorter than Beijing sublineage 2.2.2 isolates (9 SNPs, p=0.02) (**Fig. 2a**). Indeed the distribution of mean node-to-tip distances for all internal nodes was skewed significantly lower within the Beijing sublineage 2.2.1 compared to the rest of the tree (median 16 SNPs compared to 62, 57, 39 and 60 SNPs for Lineages 1, 2.1, 2.2.2 and 4, respectively; p<0.0015 in all cases). Importantly, for subtrees with the same number of descendant tips (i.e. ancestral transmission events associated with the same number of subsequently sampled *Mtb* cases), mean subtree heights were shorter within the Beijing lineage than in other lineages (**Fig. 2b**). Hence Beijing lineage subtrees represent temporally shorter transmission pathways than do subtrees of an equivalent size from other lineages. We identified potential transmission clusters using a range of maximum pairwise patristic distance thresholds to define a cluster (**Fig. 2c**). Beijing sublineage 2.2.1 was responsible for ≥70% of transmission clusters using thresholds of up to 20 SNPs (transmission age of ~20 years). Therefore, not only was sublineage 2.2.1 the most common cause of TB in the HCMC study population, but these infections resulted from more recent transmission. Coupled with the difference in age distributions (**Fig. 1**), these data indicate that new cases of Lineage 1 *Mtb* in HCMC typically result from activation of longer-term latent infections, while new cases of Lineage 2.2.1 *Mtb* often result from more recent transmission and shorter time to develop active disease.

**Figure 2.**
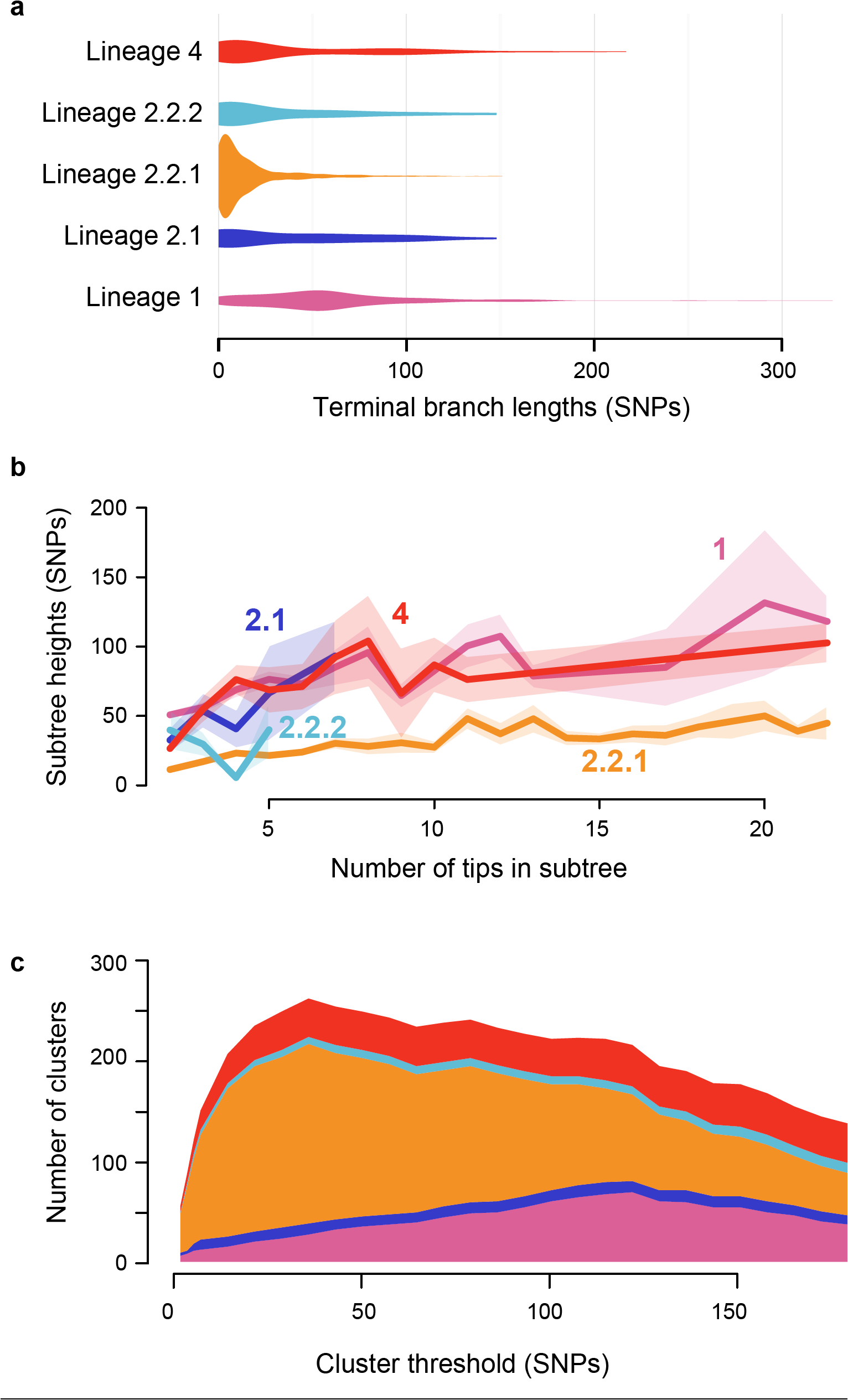
Properties of lineage subtrees for HCMC *M. tuberculosis* genomes. **(a)** Distributions of terminal branch lengths. **(b)** Mean subtree heights (y-axis; measured as mean node-to-tip distances for each subtree) vs subtree size (x-axis; number of descendant tips). Shaded region indicates standard error of the mean across subtrees of a given size; labels indicate lineage. **(c)** Stacked area plot showing number of clusters (y-axis) within each lineage (coloured as in panel a) identified using different maximum patristic distance thresholds to define clusters (x-axis).

It has been suggested previously that the Beijing lineage is slowly displacing the resident Lineage 1 strains in Vietnam, following the introduction of the Beijing strain into urban areas and subsequent spread to rural areas where Lineage 1 still dominates ^8,10^. Our data are consistent with this and suggest that the Beijing sublineage 2.2.1 isolates from HCMC may represent a locally established epidemic subclade of the Beijing lineage, similar to that previously described in Russia ^3^. To investigate this we combined our HCMC *Mtb* genome data with 3,085 publicly available *Mtb* whole genome sequences from Russia ^3^, Malawi ^4,5^, Argentina ^12^, and China ^13^, as well as globally dispersed Lineage 2 genomes ^14^ (**Supplementary Table 2**), then inferred phylogenies for Lineages 1, 2 and 4 (**Fig. 3**). Beijing sublineage 2.2.1 isolates from HCMC formed several distinct clusters that each shared a recent common ancestor with Lineage 2.2.1 isolates from outside Vietnam (**Fig. 3b**). Notably, isolates from Russia, Malawi, China and numerous other countries were interspersed throughout the HCMC sublineage 2.2.1 population (**Fig 3b**), suggesting multiple, frequent transfers of this lineage between host populations in HCMC and other geographic regions. HCMC Lineage 4 isolates were drawn from eight of the ten recognised sublineages ^6^ ‒ including those identified as specialist, generalist and intermediate in their geographic range ^15^ ‒ and were interspersed with isolates from other geographical locations, consistent with multiple imports into HCMC from external populations (**Fig. 3c**). In further support of these observations, stochastic mapping of locations onto the Lineage 2 and 4 phylogenies predicted dozens of strain transfer events between Vietnam and other locations (**Fig. 3d**) (see **Methods**). In contrast, while Lineage 1 had a similar frequency amongst Malawian isolates (16% of cases), *Mtb* isolates from the two locations were not comingled (**Fig. 3a**), suggesting that Lineage 1 associated TB in HCMC results entirely from the local endemic population.

**Figure 3.**
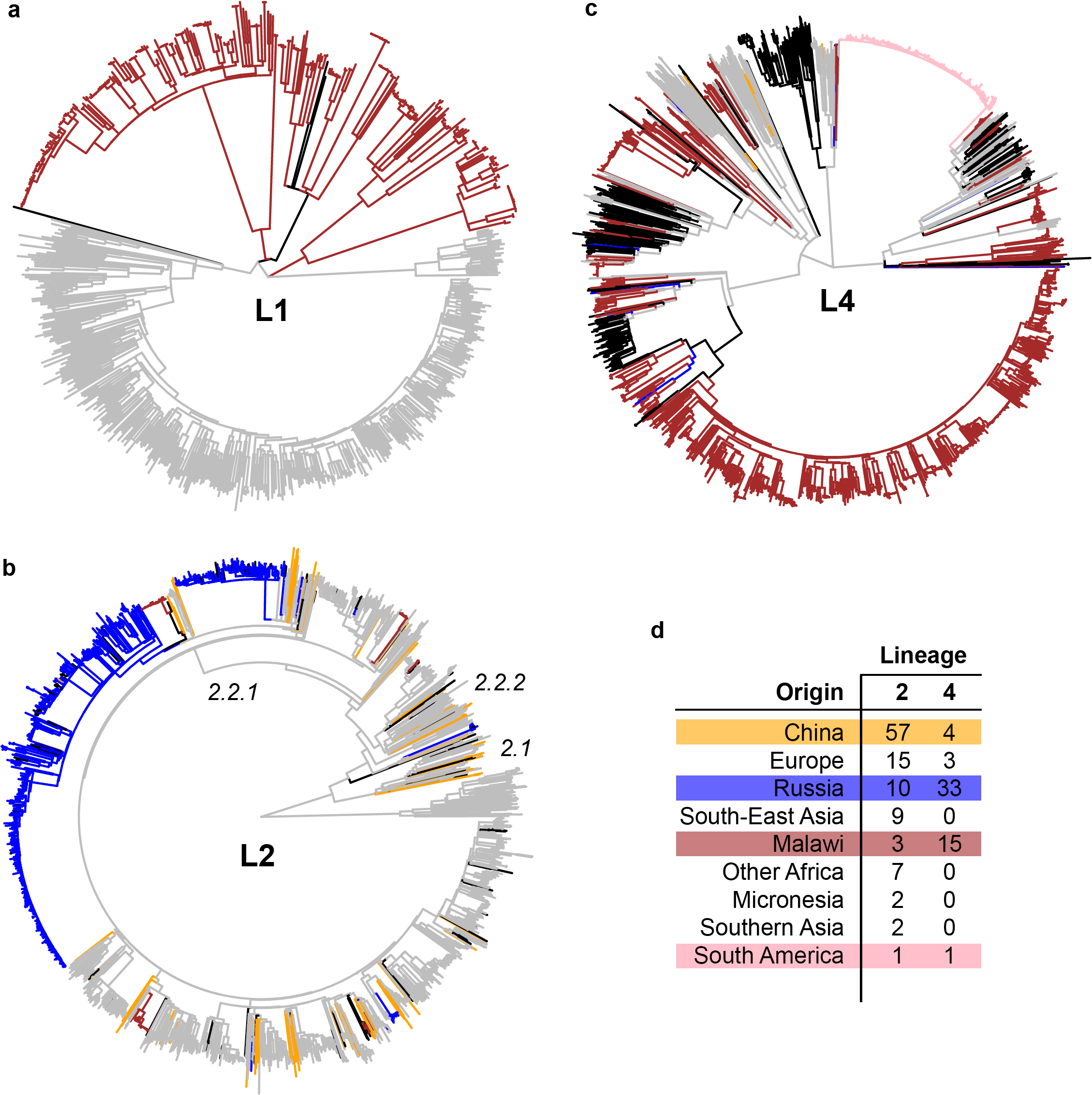
Phylogenies of *M. tuberculosis* showing relationships between isolates from HCMC and other locations. HCMC isolates are coloured grey, isolates from four other localised studies are coloured as in panel (d), other locations are shown in black. **(a)** Lineage 1 (n=675 genomes). **(b)** Lineage 2 (n=1,871 genomes). **(c)** Lineage 4 (n=2,066 genomes). **(d)** Number of transfers between Vietnam and other locations predicted by stochastic mapping of locations onto the Lineage 2 and 4 trees.

**Table 2.** Homoplasic non-synonymous SNPs identified as occurring on the Beijing lineage-defining branch and also arising independently within other lineages. The number of branches on which each SNP was identified outside the Beijing lineage-defining branch is shown, and the number of such branches that have multiple descendant tips (indicating onward transmission of the SNP) is shown in brackets. HCMC refers to the 1,635 isolates from HCMC, Vietnam; Elsewhere refers to the 3,085 additional isolates from published studies^3-5, 12-14^; trees are shown in Figure 3.

| **Mutation** | **No. branches outside Beijing lineage (no. transmitted)** |  | **Function** |
| --- | --- | --- | --- |
|  | **HCMC** | **Elsewhere** |  |
| EsxW-T2A | 9 (4) | 10 (6) | ESX-5 secreted protein<br>(CFP10 homolog) |
| Rv3081-F220L | 2 (1) | 7 (3) | hypothetical protein |
| GidB-E92D | 1 (1) | 0 (0) | streptomycin resistance |

Taken together, our data reveal a complex epidemiological landscape for TB in HCMC, comprising (i) an underlying burden of disease caused by endemic Lineage 1 *Mtb* strains (24% of all TB cases), which is associated with activation of long-term latent infection and disproportionately affects men and older people; and (ii) an additional disease burden caused by introduction of exogenous Lineage 2 and 4 *Mtb* strains (76% of all TB cases), more often affecting hosts of both genders and a broader age range and is associated with shorter time to active disease and frequent onward local transmission (>30% of all introduced strains were involved in transmission clusters defined using a maximum pairwise patristic distance threshold of ≤20 SNPs). Notably, 83% of TB cases resulting from these onward transmission events involved the Beijing sublineage 2.2.1, accounting for 23% of all cases included in the genomic study. These findings have important consequences for local TB control programs, which may benefit from distinct strategies targeting the different forms of TB that are associated with different *Mtb* lineages, and also highlight the high degree to which international transfer of *Mtb* can impact on local disease burden and shifting epidemiology.

The population structure (**Figures 1–2)** provides evidence that Beijing lineage strains are more transmissible within this HIV-negative HCMC population than are other *Mtb* lineages. Genomic evidence for enhanced transmission of the Beijing lineage has been documented in Russia (associated with antimicrobial resistance) ^3^ and Malawi (independent of antimicrobial resistance) ^4^ While antimicrobial resistance was common amongst HCMC Beijing lineage isolates, the majority of transmission clusters comprised groups of isolates that did not share any known resistance mutations that could account for their transmission success (**Supplementary Fig. 2**). This is consistent with previous reports that the Beijing lineage is highly transmissible and more likely to progress to active disease in various host populations and is also more virulent and less pro-inflammatory in various cellular assays, independent of antimicrobial resistance ^16–19^ We therefore aimed to interrogate the *Mtb* genome data to identify mutations that may contribute to the success of the Beijing lineage (2.2). Evolutionary convergence has previously been used as a signal of positive selection to identify mutations associated with antimicrobial resistance in Mtb^20,21^. We reasoned that advantageous polymorphisms contributing to the enhanced transmissibility of Lineage 2.2 should lie on the branch leading to this lineage, and should be under positive selection that is detectable as convergent evolution at these sites in other lineages. We identified a total of 420 homoplasic nonsynonymous SNPs (nsSNPs) across the HCMC phylogeny, the most frequent of which occurred in genes in which convergent evolution has previously been associated with antimicrobial resistance including *gidB, embB, gyrA, rpoB, rpoC*, and *inhA* ^20^. The homoplasic nsSNPs included three that arose on the branch defining Lineage 2.2 and also elsewhere in the HCMC tree (**Table 2**, **Supplementary Fig. 3**). One was a mutation in *EsxW* codon 2, which arose on nine other branches (six times in Lineage 4, three times in Lineage 1; see **Supplementary Fig. 3**) and showed evidence of onward transmission on four occasions. Comparison to the global tree detected the same *EsxW* mutation on a further ten Lineage 4 branches in Malawi and Russia, with onward transmission detected on six occasions. The other two mutations were in *Rv308l* (conserved hypothetical protein) and *GidB* (mutations in which are often associated with streptomycin resistance) and arose less frequently (**Table 2**). In contrast, homoplasic nsSNPs on the branches defining Lineages 1 or 4 were each detected on only one or two other branches in the HCMC tree and no additional branches of the global tree (**Supplementary Table 3**). No homoplasic SNPs were associated with sublineage 2.2.1, and although synonymous or intergenic SNPs can have functional consequences, we found no such homoplasies associated with Beijing or other lineages.

EsxW (Rv3620c) is an ESX-secreted effector protein, which has been proposed as a vaccine and immunotherapy candidate ^22–24^ due to its demonstrated immunogenicity in mice and epitopes predicted to bind a wide range of human HLA-DRB1 alleles ^22,25^. There are 23 *esx* genes in the *Mtb* genome, including 11 clustered pairs of *esx* genes whose products form secreted heterodimers. EsxW and its heterodimerization partner EsxV are encoded in adjacent genes in the RD8 locus, which is conserved in all *Mtb* genomes but lacking from other members of the *Mtb* complex ^26^. EsxW is one of five close homologs in *Mtb* (differing by 1-2 amino acids) that result from expansion of a subfamily of CFP10 homologs (QILSS/ESX-5; **Supplementary Fig. 4**). Despite their close homology, SNPs in these genes can be easily distinguished even with short reads, as the upstream sequences are unique and allow unambiguous mapping (**Supplementary Fig. 5**). The ancestral form of *Mtb* EsxW carries the polar threonine (codon ACC) at residue 2, while all other *Mtb* QILSS proteins carry the hydrophobic alanine (GCC) at this position; in the Beijing lineage EsxW residue 2 is reverted to alanine (GCC), making the protein identical to EsxJ. Residue two lies in the N-terminal loop of EsxW (**Fig. 4**), and the evolutionarily convergent changes at this position could have functional impacts such as stabilizing the heterodimer or impacting interactions with host proteins. Since the upstream promoter regions of EsxW and its homologs differ substantially (**Supplementary Fig. 5**), their expression is likely subject to different regulatory controls. While the mechanism remains to be elucidated, our results provide evidence that EsxW is under selection in natural *Mtb* populations, providing support for the prioritization of this immunomodulatory protein as a vaccine target. Importantly, our data show that genomic diversity in *Mtb* has a profound impact on TB epidemiology even within a single localized host population, and indicates that more detailed understanding of lineage-specific variation in *Mtb* could be highly informative for TB control.

**Figure 4.**
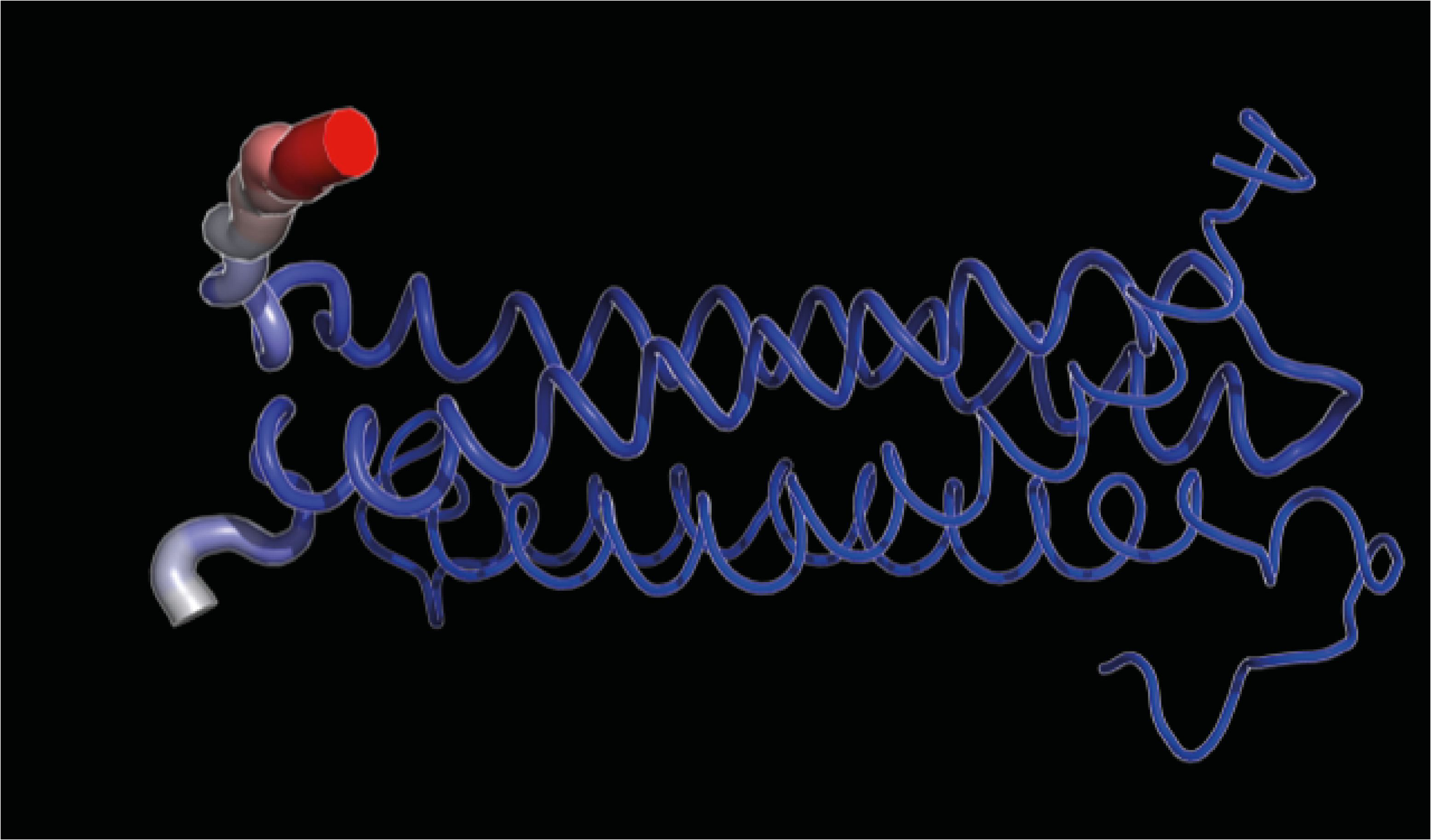
NMR structure of the *esxW* homologue *esxB* (residues 2-100). Residue 2 is highlighted in red.

## Accession codes

*Mtb* genome data was deposited in NCBI BioProject [ID:PRJNA355614; http://www.ncbi.nlm.nih.gov/bioproiect/355614]; individual accession numbers for *Mtb* genomes analysed in this study are given in **Supplementary Tables 1 and 2** (data from previous studies).

## ACKNOWLEDGMENTS

We would like to thank the clinical staff who recruited patients into our study from the following District TB Units (DTU) in HCMC, Viet Nam; District 1, 4, 5, 6, 8, Tan Binh, Binh Thanh and Phu Nhuan DTUs; and also our colleagues from Pham Ngoc Thach Hospital for Tuberculosis and Lung Disease, HCMC Viet Nam. This work was supported by the National Health and Medical Research Council, Australia (Project grant #1056689, Fellowship #1061409 to KEH, Fellowship #1061435 to MI), A*STAR Biomedical Research Council, Singapore (12/1/21/24/6689) and the Wellcome Trust UK (research training fellowship #081814/Z/06/Z to MC) and as part of their Major Overseas Program in Viet Nam (089276/Z/09/Z).

## AUTHOR CONTRIBUTIONS

SJD, KEH, MC, MI, YYT, CCK, are the study principal investigators who conceived and obtained funding for the project. SJD provided overall project co-ordination; MI organized and supervised the DNA sequencing and KEH devised the overall analysis plan and wrote the first draft of the manuscript along with PM. MC and SJD established the TB cohort for this genetics study by working with PVKT, DTMH, NNL, NHL, NTQN, NTTT, GT and JJF to coordinate the collection of clinical samples and phenotypes. KP performed DNA quality checks and sequencing on all Vietnamese samples. KEH, PM, MI, DJE analyzed the data. All authors critically reviewed manuscript revisions and contributed intellectual input to the final submission.

## Online Methods

**Bacterial isolates used in this study.** Between December 2008 and July 2011, 2,091 individuals of the Vietnamese Kinh ethnic group attending the outpatient department of Pham Ngoc Thach Hospital or from 8 District Tuberculosis Units (District 1, 4, 5, 6, 8, Tan Binh, Binh Thanh and Phu Nhuan) in HCMC were recruited into a clinical study of TB. Consenting adult patients were recruited on the basis of: (1) sputum smear positivity, (2) no evidence of HIV infection, and (3) no prior history of TB antibiotic therapy. *Mtb* strains were isolated from the study participants, resulting in a culture collection of N=1822 *Mtb* isolates.

**DNA extraction and sequencing.** *Mtb* isolates were subcultured on Lowenstein Jensen media and DNA extracted at the Oxford University Clinical Research Unit in HCMC using the cetyl trimethylammonium bromide (CTAB) extraction protocol as described previously ^27^ DNA was successfully obtained from N=1,728 isolates and shipped to the University of Melbourne for whole genome sequencing. DNA extracts were purified using AxyPrep™ Mag PCR Normalizer Protocol prior to library preparation. A total of N=1,655 DNA samples passed QC and were subjected to library preparation using the Nextera XT protocol. Libraries were quantified using Quant-iT PicoGreen (dsDNA kit, Invitrogen), then normalised and pooled to 4 nM concentration. DNA underwent 150 bp paired end sequencing (Rapid mode v2) on the Illumina HiSeq 2500 platform (Illumina, San Diego). Sequence data was successfully generated for N=1,636 *Mtb* isolates from HCMC (representing 90% of those isolated from eligible patients in the cohort) with median three million reads per sample, providing median 99.2% coverage and 86x depth for each *Mtb* genome (**Supplementary Table 1**).

**Publicly available genome data used in this study.** Illumina *Mtb* genome sequences from various previously published studies were downloaded from the European Nucleotide Archive (accessions in **Supplementary Table 2**). A total of 3,085 *Mtb* genomes were included in the analysis, comprising data from localized studies: 1,032 from Russia ^3^, 1,621 from Malawi ^4,5^, 248 Argentina ^12^, and 78 from China ^13^; as well as 106 globally dispersed Lineage 2 genomes ^14^. The H37Rv reference genome sequence (accession NC_000962.3) was used for all reference-driven analyses.

**SNP analysis.** Sequence reads were mapped to the H37Rv reference genome using the RedDog pipeline v0.5 (https://github.com/katholt/RedDog). Briefly, Bowtie2 v2.2.3 was used for read alignment with the sensitive-local algorithm and the maximum insert length set to 2000 (via the -x parameter) ^28^ and variant sites (SNPs) were called using SAMTools v0.1.19 ^29^ SNPs located in previously reported repetitive regions of the genome were excluded prior to phylogenetic analysis ^30,31^ (**Supplementary Table 4**); sites for which a definitive allele call could not be made in at least 99.5% of all isolate sequences were also excluded from the set of SNPs used for phylogenetic analysis. Two SNP alignments were compiled for analysis: one comprising the 1,635 HCMC isolates (total 73,718 SNPs), and one comprising all 4,720 isolates (including the HCMC isolates and the global collections downloaded from public data; total 133,495 SNPs).

***In silico* lineage and antimicrobial resistance typing.** Mykrobe Predictor was used to analyse raw Illumina reads generated from HCMC *Mtb* isolates and (a) assign each isolate to one of the seven *Mtb* lineages, and (b) detect known resistance associated polymorphisms ^32^ (summarized in **Table 1,** individual mutation calls are provided in **Supplementary Table 1**). All *Mtb* isolates were further assigned to sublineages by comparing SNPs identified using RedDog with those used in the haplotyping scheme defined by Coll *et al*. ^6^ (lineage assignments are in **Supplementary Tables 1-2**).

**Phylogenomic analyses.** ML phylogenetic trees were inferred using RAxML v7.7.2 ^33^ for (a) all HCMC isolates (presented in **Fig. 1**); and (b) each of lineages 1, 2 and 4 using combined data from the HCMC isolates and available public data (presented in **Fig. 3;** see isolates list in **Supplementary Tables 1-2**). The trees presented are those with the highest likelihood from 5 replicate runs, constructed using the GTR model of nucleotide substitution and a Gamma model of rate heterogeneity to analyse a concatenated alignment of SNP alleles. An approximate ML tree containing all data (HCMC isolates and available public data) was inferred using FastTree v2.1.8 ^34^ Ancestral sequence reconstruction was performed for the HCMC tree and combined tree using FastML v3.1 to infer the sequence alignment at each internal node of the ML phylogeny ^35^. Substitution events occurring on each branch of the tree were extracted by comparing the joint reconstruction sequences for the parent and child nodes; these data were used to identify lineage-specific polymorphisms and to detect independent occurrences of those polymorphisms outside of the lineage of interest (data in **Table 2**). Terminal branch lengths reported are the number of substitutions (SNPs) mapped to each terminal branch (data in **Fig. 2a**). Metrics for genetic diversity and tree topology were calculated from the phylogenies using R (see **Supplementary Fig. 1**). Mean subtree heights were defined as the mean root-to-tip distance for all tips in the subtree; the width of a subtree was defined as the number of descendant tips (data in **Fig. 2b**). Clusters were defined as subtrees for whom the maximum patristic distance between descendant tips (see **Supplementary Fig. 1**) fell below a specified threshold (data in **Fig. 2c**). Each cluster was checked to determine whether all members of the cluster shared any of the antimicrobial resistance mutations identified by Mykrobe Predictor; clusters in which no known antimicrobial resistance mutation was conserved in all members of the cluster are reported as not explained by antimicrobial resistance (data in **Supplementary Fig. 2**).

**Phylogeography analysis.** Transmission between geographical regions was assessed separately for the Lineage 2 and Lineage 4 trees using an implementation of stochastic mapping on phylogenies (SIMMAP) implemented in the phytools v0.5 package for R ^36,37^ Region of origin was treated as a discrete trait and mapped to each tree using the ARD model (which allows each region-to-region transfer rate to vary independently) with 100 replicates. The results reported (**Fig. 3d**) are the median values for the number of transitions to Vietnam from any other region, summarized from 100 replicate mappings for each tree.

**Esx sequence analysis.** Esx protein sequences were extracted from the H37Rv reference genome using Artemis, aligned using Muscle, and subjected to phylogenetic inference using PhyML (**Supplementary Fig. 4**). DNA sequences (length 500 bp) upstream of each *esx* gene were extracted from the H37Rv reference genome using Artemis and aligned and visualised using JalView (**Supplementary Fig. 5**).

